# The Impact of Edema and Fiber Crossing on Diffusion MRI Metrics: DBSI vs. Diffusion ODF

**DOI:** 10.1101/821082

**Authors:** Zezhong Ye, Sam E. Gary, Peng Sun, Sourajit Mitra Mustafi, George Russell Glenn, Fang-Cheng Yeh, Harri Merisaari, Guo-Shu Huang, Hung-Wen Kao, Chien-Yuan Lin, Yu-Chien Wu, Jens H. Jensen, Sheng-Kwei Song

**Affiliations:** Department of Radiology, Washington University School of Medicine, St. Louis, MO; Medical Scientist Training Program, University of Alabama at Birmingham, Birmingham, AL; Department of Radiology and Imaging Sciences, Indiana University School of Medicine, Indianapolis, IN; Department of Radiology and Imaging Science, Emory University, Atlanta, GA; Department of Neurological Surgery, University of Pittsburgh, Pittsburgh, PA; Department of Diagnostic Radiology, University of Turku, Turku, Finland; Department of Radiology, Tri-Service General Hospital, National Defense Medical Center, Taipei, Taiwan; GE Healthcare, Taipei, Taiwan; Department of Radiology and Radiological Science, Medical University of South Carolina, Charleston, SC; Department of Neuroscience, Medical University of South Carolina, Charleston, SC; Center for Biomedical Imaging, Medical University of South Carolina, Charleston, SC

**Keywords:** brain tumor, diffusion basis spectrum imaging, diffusion tensor imaging, diffusion kurtosis imaging, generalized q-sampling imaging, neurite orientation dispersion and density imaging, q-ball imaging, white matter tractography

## Abstract

**Purpose:** Diffusion tensor imaging (DTI) has been employed for over two decades to noninvasively quantify central nervous system (CNS) diseases/injuries. However, DTI is an inadequate simplification of diffusion modeling in the presence of co-existing inflammation, edema, and crossing nerve fibers.

**Methods:** We employed a tissue phantom using fixed mouse trigeminal nerves coated with various amounts of agarose gel to mimic crossing fibers in the presence of vasogenic edema. Diffusivity measures derived by DTI and diffusion basis spectrum imaging (DBSI) were compared at increasing levels of simulated edema and degrees of fiber crossing. Further, we assessed the ability of DBSI, diffusion kurtosis imaging (DKI), generalized q-sampling imaging (GQI), q-ball imaging (QBI), and neurite orientation dispersion and density imaging (NODDI) to resolve fiber crossing, in reference to the gold standard angles measured from structural images.

**Results:** DTI-computed diffusivities and fractional anisotropy (FA) were significantly confounded by gel-mimicked edema and crossing fibers. Conversely, DBSI calculated accurate diffusivities of individual fibers regardless of the extent of simulated edema and degrees of fiber crossing angles. Additionaly, DBSI accurately and consistently estimated crossing angles in various conditions of gel-mimicked edema when comparing with gold standard (r^2^=0.92, p=1.9×10^−9^, bias=3.9°). Small crossing angles and edema sinficantly impact dODF, making DKI, GQI and QBI less accurate in detecting and estimating fibers corrsing angles. Lastly, we demonstrate DBSI’s superiority over DTI for recovering and delineating white matter tracts in peritumoral edema for preoperative planning of surgical resection.

**Conclusions:** DBSI is able to separate two crossing fibers and accurately recover their diffusivities in a complex environment characterized by increasing crossing angles and amounts of gel-mimicked edema. DBSI also indicated better angular resolution capability compared with DKI, QBI and GQI.

## 1. INTRODUCTION

Diffusion tensor imaging (DTI) does not accurately estimate diffusion parameters in the presence of extra-fiber pathology, e.g., edema and cell infiltration. DTI is also unable to provide accurate neural architecture information in the presence of crossing fibers. Multiple advanced diffusion MRI methods have been proposed to better extract the non-Gaussian diffusion-weighted data that DTI fails to incorporate into its analyses.

Q-space diffusion imaging, e.g., diffusion spectrum imaging (DSI), estimates the diffusion orientation distribution function (dODF),^1^ which describes the probability of water diffusion within a voxel along specific directions. The dODF is commonly used for fiber tracking and connectivity analyses.^2^ DSI requires large pulsed field gradients and time-intensive sampling on a three-dimensional Cartesian lattice.^3–5^ High angular resolution diffusion imaging (HARDI) resolves intravoxel fiber crossing via sampling of spherical shells rather than three-dimensional Cartesian planes.^6^ One variant of HARDI, q-ball imaging (QBI), utilizes a Funk-Radon transform, reconstructing the HARDI signal to resolve multiple intravoxel fiber crossings in both cortical and deep subcortical white matter pathways without assuming Gaussian diffusion.^5,7^ QBI samples and creates the dODF via a single HARDI shell, thereby eliminating Cartesian construction bias.

Generalized q-sampling imaging (GQI) can model diffusion from the shell sampling scheme used in QBI or the grid sampling scheme used in DSI.^8^ However, GQI is unlike QBI and DSI in that GQI calculates the spin distribution function (SDF) rather than the dODF, thus directly quantifying distribution of the spins that undergo diffusion rather than dODF’s estimation of the probability of diffusion displacement.^8^ Another fundamental difference between SDF and dODF is SDF includes spin density of protons, which provides commonality between spins and enables comparison of SDFs across individual voxels.^8^

Diffusion kurtosis imaging (DKI) is a clinically feasible diffusion MRI method that employs low b-value data (maximum b-value of about 2000 s/mm^2^), maintains a short scan time, and provides an adequate signal-to-noise ratio (SNR).^9,10,11^ As with DTI, DKI robustly quantifies several physical properties of water diffusion, but is more comprehensive than DTI in characterizing diffusional non-Gaussianity. DKI can be augmented with tissue models to help interpret the biological meaning of diffusional changes caused by disease.^12,13^ In contrast to DTI, dODFs calculated with DKI are able to resolve fiber crossings.^13,14^ Tractography generated with DKI dODFs has been shown to be comparable to DSI tractography.^15^

Lastly, neurite orientation dispersion and density imaging (NODDI) utilizes a three-tissue compartment (intracellular, extracellular, and CSF, respectively) to model diffusion-weighted signals using a two-shell HARDI data acquisition scheme.^16^ To estimate neural connectivity and orientation, NODDI derives the orientation dispersion index (ODI), which summarizes the angular variation of neurites.^16^ However, NODDI does not provide estimates for fiber crossing angles.

Our lab has developed diffusion basis spectrum imaging (DBSI), which employs a unique multitensor approach that models diffusion characteristics of individual image voxels independent of a predetermined tissue model.^17–19^ DBSI assumes no exchange between intra- and extra-axonal compartments. It models diffusion-weighted MRI signals using a linear combination of multiple tensors.^17^ DBSI differs from DTI and other models in that it separates isotropic diffusion components to discriminate between inflammation, cellularity, and cerebrospinal fluid (CSF), enabling tracking of pathologies such as multiple sclerosis,^18,20,21^ cervical spondylotic myelopathy,^22,23^ traumatic spinal cord injury,^24,25^ epilepsy,^26^ and intracranial inflammation in HIV+ patients.^27^ Importantly, DBSI simultaneously resolves angle of crossing fibers and quantifies individual fiber diffusivity, a feature that neither DTI, DKI, QBI nor NODDI possesses.

Given the previous DBSI findings, the goal of this study was to validate DBSI’s ability to estimate individual nerve diffusivity, as well as compare its and other advanced diffusion MRI techniques’ ability to estimate the angle of crossing fibers in a simulated edematous environment. We compared the DBSI-derived axial diffusivity (AD) and radial diffusivity (RD) results to DTI-derived AD, RD, and fractional anisotropy (FA). Additionally, we compared DBSI’s crossing angle estimations with leading dODF methods like QBI, GQI, and DKI. Our results have widespread clinical implications for implementing DBSI as an alternative to DTI for quantifying nerve diffusivities in edematous disease states and as an alternative to DTI and dODF models for fiber tracking and preoperative surgical planning.

## 2 MATERIALS AND METHODS

All experimental procedures involving animals were approved by Washington University’s Animal Studies Committee and conformed to the Public Health Service Policy on Humane Care and Use of Laboratory Animals (http://grants.nih.gov/grants/olaw/olaw.htm).

### 2.1 Preparation of fixed mouse trigeminal nerve phantoms

Five female C57BL/6 mice (The Jackson Laboratory) of 8-12 weeks of age were euthanized and perfused with 1% phosphate buffered saline (PBS) solution and fixed in 4% paraformaldehyde (PFA) solution for 24 hours. After fixation, a total of ten trigeminal nerves were extracted from the fixed mice and placed in 1% PBS for 48 hours. Individual nerves were scanned *ex vivo* to acquire their respective baseline diffusion properties. To simulate crossing fibers, two nerves were juxtaposed tightly in parallel, aligned with a single nerve, and scanned at three different crossing angles: 90°, 60°, and 40°, respectively. Vasogenic edema was simulated by coating the single nerve and the juxtaposed nerves with 4% agarose gel. Increased amounts of edema were further simulated by adding more gel between the single fiber and the juxtaposed fibers (Figure 1). Nerve combinations were scanned with the same imaging parameters as the baseline individual nerve scans, as described below.

**Figure 1.**
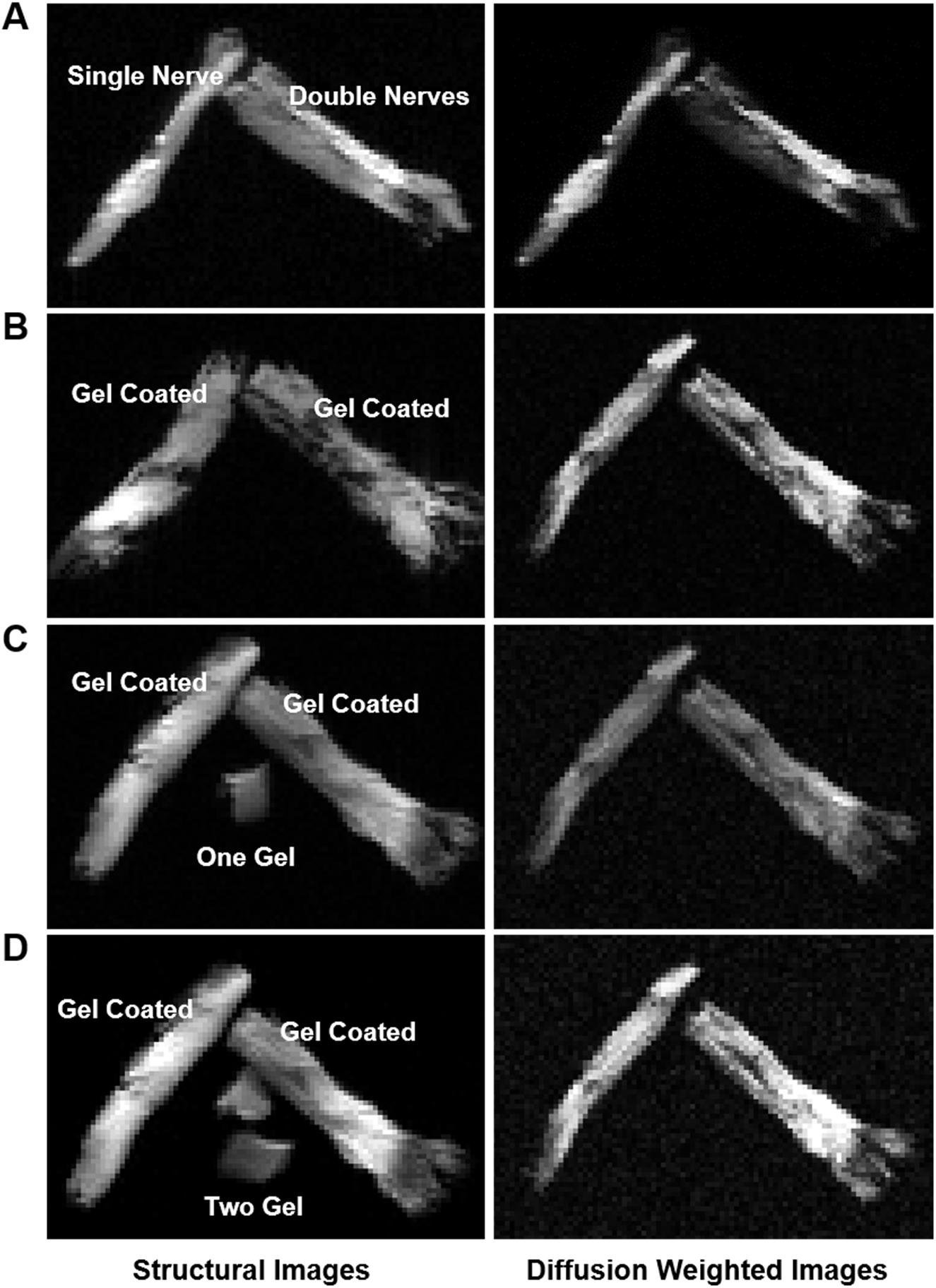
Structural images and diffusion weighted images of trigeminal nerves. Nerves were placed in a 1:2 ratio: single nerve (single nerve) and two different nerves juxtaposed in parallel (double nerves). (A) 90° without gel; (B) 90° with gel coating; (C) 90° with gel coating and one additional piece of gel; (D) 90° with gel coating and two pieces of gel. Gel consisted of 4% agarose. The same procedure was followed for scans of 60° and 40°.

### 2.2 Diffusion-weighted spectroscopy of fixed trigeminal nerve phantoms

MR imaging and spectroscopy were performed using a 4.7T MR scanner (console by Agilent Technologies, Santa Clara, CA; magnet by Oxford Instruments, Oxford, UK; gradients by Magnex Scientific, Oxford, UK), and an actively shielded Magnex gradient coil (60 G/cm, 270 µs rise time). A home-made surface coil with inner diameter of 1 cm was built specifically for trigeminal nerve imaging. The diffusion-weighted spectroscopy parameters were as follows: TR=1500 ms, TE=40 ms, time between application of gradient pulse (∆)=25 ms, diffusion gradient time (δ)=8 ms, and number of average=1. A multi-echo spin-echo diffusion-weighted sequence was employed for data acquisition using diffusion weighting schemes described in the following methods: DBSI (99 diffusion encoding directions, max b-value=3000 s/mm^2^);^17^ Linearly Estimated Moments provide Orientations of Neurites And their Diffusivities Exactly (LEMONADE: 325 diffusion encoding directions, max b-value=4000 s/mm^2^);^28^ NODDI (145 diffusion encoding directions, max bvalue=4000 s/mm^2^);^16^ and hybrid diffusion imaging (HYDI: 143 diffusion encoding directions, max b-value=6000 s/mm^2^).^29^

### 2.3 Structural MRI of fixed trigeminal nerve phantoms

Structural imaging and diffusion-weighted imaging (DWI) was performed using the same MR scanner and surface coil. Structural images were acquired with a spin-echo sequence: TR=600 ms, TE=20 ms, flip angle=30°, field of view=24×24 mm^2^, matrix size=128×128, and slice thickness=1 mm. Diffusion-weighted images were acquired using a diffusion-weighted spin-echo sequence: TR=1500 ms, TE=40 ms, b-value=1500 s/mm^2^, field of view=24×24 mm^2^, matrix size=128×128, and slice thickness=1 mm.

### 2.4 *In Vivo* MRI of human subject

One patient with metastatic lung adenocarcinoma underwent imaging on a GE Discovery MR 750 3T MRI scanner (GE Healthcare, Waukesha, WI, USA). The study was approved by the local Institutional Review Board, and informed consent was obtained and documented from the participant. Diffusion-weighted images were acquired using a 99-direction diffusion-weighting encoding scheme (maximum b-value=1500 s/mm^2^) using an echo planar imaging (EPI) sequence: TR=6000 ms, TE=88 ms, FOV=256×256 mm^2^, slice thickness=2.5 mm, in-plane resolution=0.94×0.94 mm^3^, and total acquisition time=19 min.

### 2.5 Diffusion MR data processing and analyses

#### 2.5.1 DTI analyses

Diffusion-weighted data with 99 diffusion directions was analyzed with the DTI single-tensor model analysis package developed in-house with Matlab Version 2013 (MathWorks; Natick, MA, USA) software.

#### 2.5.2 DBSI analyses

Diffusion-weighted data with 99 diffusion directions was analyzed with the DBSI multi-tensor model analysis package developed in-house with Matlab Version 2013 software. As detailed in Wang, et al.,^17^ DBSI models the diffusion-weighted MR signal as a linear combination of discrete anisotropic diffusion tensors (first term in Eq. [1]; reflecting nerve fibers) and a spectrum of isotropic diffusion tensors (second term; reflecting inflammatory cells, edema, and CSF).

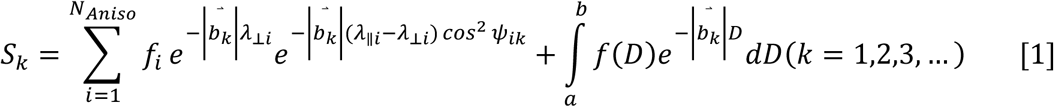

The quantities *S*_*k*_ and 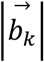 are the normalized signal and *b*-value of the *k*^*th*^ diffusion gradient; *N*_*Aniso*_ is the number of anisotropic tensors to be determined; *λ*_||*i*_ and *λ*_⊥*i*_ are the axial and radial diffusivities of the *i*^*th*^ anisotropic tensor under the assumption of cylindrical symmetry; ψ_*ik*_ is the angle between the diffusion gradient *b*_*k*_ and the principal direction of the *i*^*th*^ anisotropic tensor; *f*_*i*_ is the signal intensity fraction of individual anisotropic tensor components; and *a* and *b* are the low and high diffusivity limits for the isotropic diffusion spectrum *f*(*D*).

Anisotropic diffusion tensors were defined to detect different fibers. Coordinates of two different anisotropic tensors could be used to resolve crossing fibers and calculate their crossing angles. Anisotropic tensor signal intensity fractions (fiber fractions) were used to separate two different fibers. AD and RD were used to assess directional diffusivity of the fibers. An isotropic diffusion spectrum was derived and analyzed to define tensors for cell infiltration, vasogenic edema, and cerebrospinal fluid (CSF).

#### 2.5.3 DKI analyses

Diffusion kurtosis imaging (DKI) provides a general and rigorous means of quantifying diffusional non-Gaussianity in complex diffusive media, such as biological tissues.^9^ Jensen and colleagues derived the DKI-derived dODF, or kurtosis dODF [Eq. 2], as a corrected version of the Gaussian ODF so as to account for non-Gaussian diffusion.^9,10^

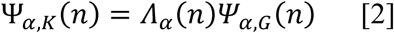

where Ψ_*a,K*_(*n*) and *ψ*_*a,G*_(*n*) refer to the kurtosis dODF and Gaussian dODF approximations, respectively. Λ_*α*_(*n*) is a correction factor that accounts for non-Gaussian diffusion. Maxima of the kurtosis dODF indicate directions of less hindered diffusion and in white matter are typically assumed to approximate the orientations of axonal fiber bundles. In contrast to the Gaussian dODF, which only identifies a single orientation per voxel, the kurtosis dODF is able to detect multiple orientations and thereby resolve fiber crossings. Nonetheless, the kurtosis dODF, as with other dODFs, has a limited resolving power and so may fail to detect or accurately quantify fiber crossings with smaller angles.^30^ DKI dODF reconstruction were performed with alpha=4 on the LEMONADE dataset at b=0, 1500, and 3000 s/mm^2^ using the methods described by Glenn and coworkers.^9^

#### 2.5.4 GQI analyses

GQI calculates the spin distribution function (SDF; Eq. [3]), which represents the distribution of the spins of water molecules as they diffuse.^8^

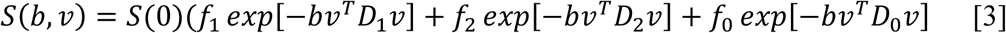

where b and *ν* represent the b-value and unit vector of the diffusion gradient, respectively, *f*_1_ and *f*_2_ are the volume fractions of the two fiber bundles, and *f*_0_ is the volume fraction of isotropic diffusion. *D*_1_, *D*_2_, and *D*_0_ are the diffusion tensors for the three corresponding volume fractions. The fiber orientations were determined by the local maxima of the reconstructed SDFs. The fiber crossing angles were evaluated by resolving major and minor fibers. The major fiber and minor fiber were defined by the largest local maximum (the global maximum) and the second largest local maximum, respectively. We used GQI in this study to generate SDFs using the LEMONADE dataset.

#### 2.5.5 NODDI analyses

In neurite orientation dispersion and density imaging (NODDI) introduced by Zhang et al.,^16^ the tissue is modelled with three compartments as:

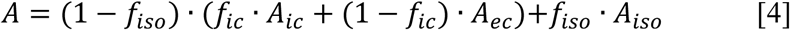

where *f*_*ic*_, *f*_*ec*_, and *f*_*iso*_ are the volume fractions for intra-cellular, hindered extra-cellular and isotropic CSF components, and *A*_*ic*_, *A*_*ec*_, and *A*_*iso*_ are the corresponding normalized signals. The orientation dispersion index (ODI) is the summary statistic of NODDI and is calculated from concentration parameter *κ*, which measures the extent of orientation dispersion about the mean neurite orientation. ODI is defined as:

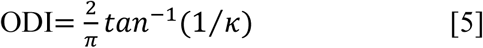

NODDI results were obtained by fitting data using the NODDI toolbox (NODDI toolbox v0.9., http://www.nitrc.org/projects/noddi_toolbox/) with CSF volume fraction diffusivity fixed at 2.0 *µm*^2^/*ms*. A Matlab code was developed to fit signals into a hierarchy of two stages of multiple initializations to avoid local minima. The initialization values for the first step were: *f*_*ic*_ = {0, …, 1} (step size 0.1), *A*_*ic*_ = {0.5, …, 1.8} *µm*^2^/*ms*, (step size 0.1 *µm*^2^/*ms*), *f*_*iso*_ = {0.1, …, 0.3} (step size 0.1), *κ* = {0.0, …, 0.5}, (step size 0.1). The second step had initializations between upper and lower bounds as nearest initialization values around the found minima in the first stage. Step sizes for the second stage of multiple initializations were 0.01 for *f*_*ic*_, *A*_*ic*_, *f*_*iso*_ and *κ*,. The *A*_*ic*_ was initialized in the NODDI toolbox as: *A*_*ic*_ = *A*_*ic*_ ∗ (1 − *f*_*ic*_), and the mean neurite orientation by fitting single tensor as in DTI.

### 2.5.6 QBI analyses

Tuch, et al.,^7^ presented the QBI-derived ODF as follows:

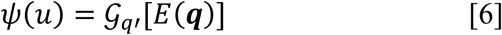

*where ψ*(*u*) is the ODF, 𝒢_*q′*_, is the Funk-Radon transform, q׳ is the radius of the sampling shell, and **q** is the diffusion wavevector.^7^ Specifically, the relationship between the QBI-derived ODF and the Funk-Radon transform is:

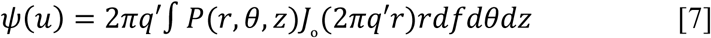

Where P(r,θ,z) is the PDF in cylindrical coordinates and J_ₒ_ is the zeroth-order Bessel function. By summing the diffusion signal along the equator around a particular direction, the diffusion probability in that direction could be estimated. This provides a model-free approach for estimating the diffusion probability from the spherically sampled diffusion signal. QBI analyses were performed on the HYDI dataset. Maximum likelihood estimation was used to estimate the fiber orientation from the ODF.

### 2.6 Calculation of gold standard fiber crossing angles

The gold standard fiber crossing angles were calculated from each corresponding DW image, analyzed using ImageJ (NIH, Bethesda, MD). Based on the DWI (Fig. 2A), binary masks were generated (Fig. 2B), and linear regression analysis was performed individually on each fiber mask (Fig. 2B, red lines). We then calculated the crossing angle from the linear regression as the gold standard for diffusion MR-derived angle comparisons (Fig. 2C, θ).

**Figure 2.**
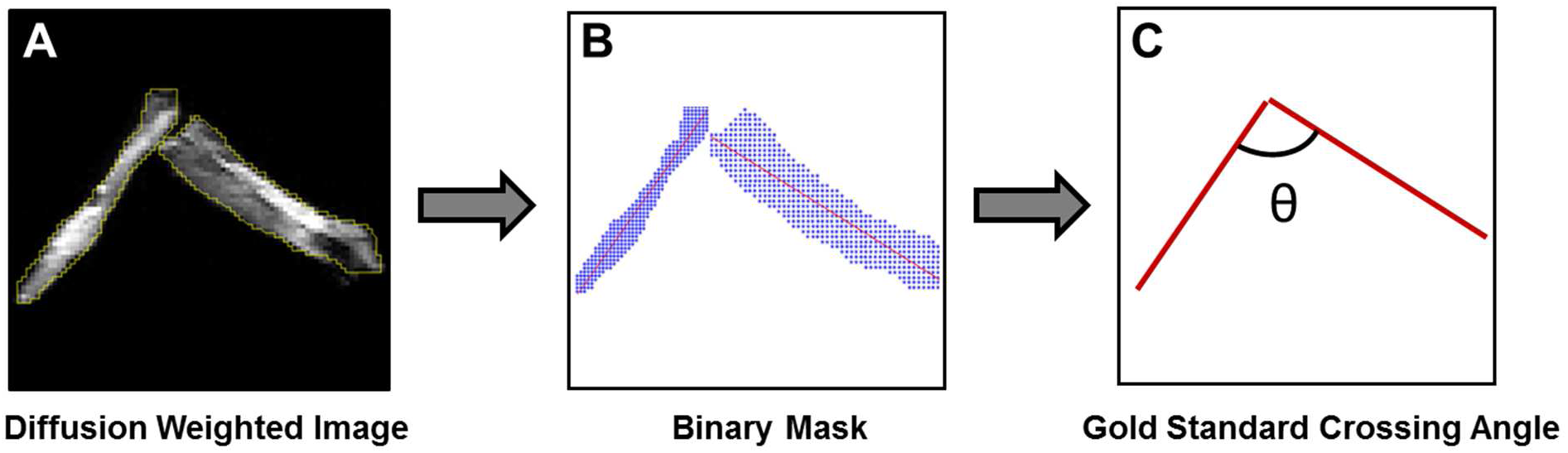
Fiber crossing angles were calculated from diffusion weighted images (A) via linear regression analysis (B). (A) Regions of interest (ROI) drawn on the diffusion weighted image. (B) Binary mask and linear regression analyses were applied to estimate the crossing angle. This crossing angle served as the benchmark for MR-derived angle calculations.

### 2.7 DBSI and DTI whole brain tractography of metastatic brain tumor patient

Modified whole brain streamlined fiber tracking^1^ was conducted using the fiber orientations derived by DBSI. Starting locations of the tracts were randomly placed within the whole brain or selected region of interest. If more than one fiber was identified by DBSI, the initial direction was randomly chosen from the resolved crossing fibers. Trilinear interpolation was used to estimate the propagation direction. The step size was 0.5mm (half of the spacing), and the maximum turning angle was 60°. DBSI-derived fiber fraction (≥15%) was used as the threshold to define a fiber. DTI used FA threshold to define a fiber (FA>0.2). DTI-derived and DBSI-derived fiber orientation was color-coded using DSI Studio (http://dsi-studio.labsolver.org).^31^ Solid tumor was defined and rendered based on gadolinium-enhanced T1W images. ROIs were drawn on all imaging slices that contained gadolinium enhancements. Vasogenic edema was defined and rendered from fluid attenuated inversion recovery images (FLAIR). Both solid tumor and vasogenic edema renderings were performed by DSI Studio.

### 2.8 Statistical analysis

Two samples student’s t-test was used to compare two different test groups. A difference of p<0.05 was considered significant. For multiple group comparisons, Bonferroni correction was used to adjust the p values. Pearson’s correlation was used to measure strengths of monotonic increasing or decreasing association between diffusion MRI estimated crossing angles (or ODI) and gold standard angles. Bland-Altman analysis was also used to evaluate the agreement between DBSI-estimated crossing angles and gold standard angles.

## 3 RESULTS

### 3.1 DTI failed to accurately resolve crossing angles, AD, RD, and FA

DTI modeled crossing fibers with single tensors and resulted in decreased AD and increased RD values compared with single nerve baseline. As crossing angles increased, the AD values decreased (Fig. 3A, single nerve: 0.77±0.04 µm^2^/ms; 40°: 0.67±0.04 µm^2^/ms; 60°: 0.65±0.07 µm^2^/ms; 90°: 0.60±0.06 µm^2^/ms), and the RD values increased (Fig. 3B, single nerve: 0.15±0.04 µm^2^/ms; 40°: 0.14±0 µm^2^/ms; 60°: 0.27±0.06 µm^2^/ms; 90°: 0.33±0.07 µm^2^/ms). Compared to the single nerve baseline, these AD and RD values were significantly different (p<0.05), except for the RD values at 40°. These systemic errors complicate evaluations of axon and myelin integrity using DTI AD and RD in the presence of fiber crossing. As a result of the decreased AD and increased RD, the DTI FA decreased in edematous environments (Fig. 3C). The FA values for single nerve baseline, 40° crossing angle, 60° crossing angle and 90° crossing angle were 0.77±0.06, 0.75±0.01, 0.48±0.06 and 0.36±0.10, respectively. Compared with baseline, there were statistically significant differences for 60° crossing angle (p<0.05) and 90° crossing angle (p<0.05), respectively. We also compared the AD, RD and FA values between crossing fibers with gel and without gel. Compared with crossing fibers without gel, both AD values (60°: 0.61±0.06 µm^2^/ms vs. 0.66±0.07 µm^2^/ms, p=0.39; 90°: 0.55±0.02 µm^2^/ms vs. 0.62±0.06 µm^2^/ms, p=0.01; Fig. 3D) and RD values (60°: 0.22±0.03 µm^2^/ms vs. 0.29±0.06 µm^2^/ms, p=0.036; 90°: 0.24±0.04 µm^2^/ms vs. 0.37±0.03 µm^2^/ms, p=0.0057; Fig. 3E) of crossing fibers with gel increased. FA values decreased in fibers with gel relative to the same fibers without gel at 60 and 90 degrees of fiber crossing. (60°: 0.57±0.05 vs. 0.48±0.06, p=0.04; 90°: 0.47±0.09 vs. 0.30±0.04, p=0.002; Fig. 3F).

**Figure 3.**
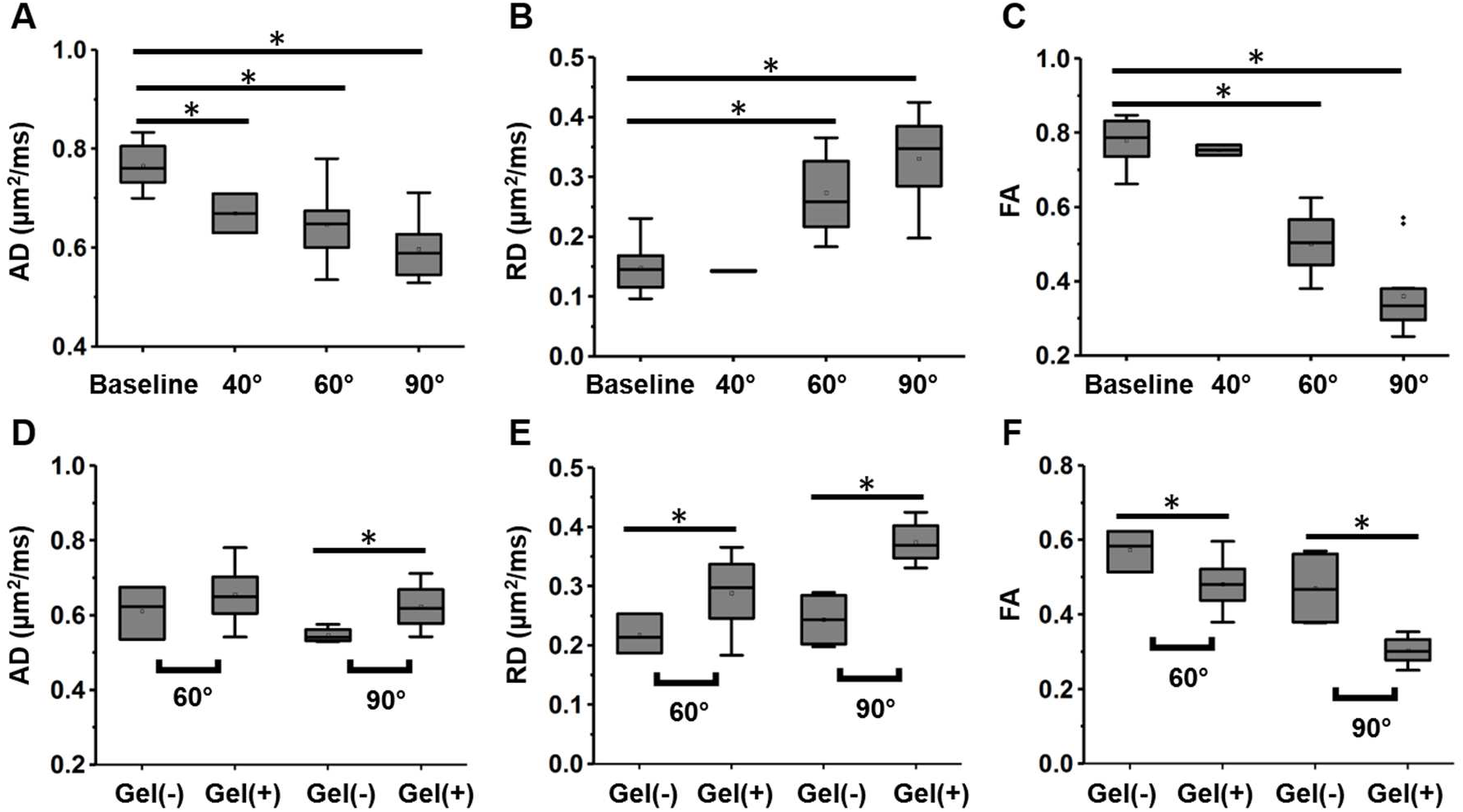
Comparison of DTI-derived diffusivity at three fiber crossing angles, as well as environments mimicking healthy (gel-) and edematous (gel+) environments. DTI modeled decreased AD (A) and increased RD (B) in comparison to single nerve baseline as crossing angles increased. DTI FA decreased as the angle of crossing fibers increased (C). Gel-added crossing fibers for both 60° and 90° show increased AD (D), RD (E) and decreased FA (F) compared to crossing fibers without gel. P values were adjusted suing Bonferroni correction. *\*P*<0.05.

### 3.2 DBSI accurately identified crossing fibers, quantified their diffusivities, and resolved crossing angles under various edematous conditions

In contrast to DTI, DBSI resolved crossing fibers and quantified individual fiber fractions, with fiber-1 (two nerves juxtaposed) fiber fraction > fiber-2 fiber fraction (single nerve) (without gel: 0.56±0.08 vs. 0.31±0.05, p=6.9×10^−13^; with gel: 0.51±0.08 vs. 0.23±0.09, p=1.0×10^−5^; Fig. 4A). To validate the DBSI-estimated fiber fraction, fiber volumes were calculated from corresponding diffusion-weighted images as gold standard. Gel-mimicked edema was also detected and quantified by DBSI hindered fraction. The DBSI hindered fraction for crossing fibers with gel were significantly higher than those from single nerve baseline (0.19±0.3 vs. 0.03±0.04, p=4.4×10^−9^) and crossing fibers without gel (0.19±0.3 vs. 0.04±0.04, p=4.6×10^−9^) (Fig. 4C).

**Figure 4.**
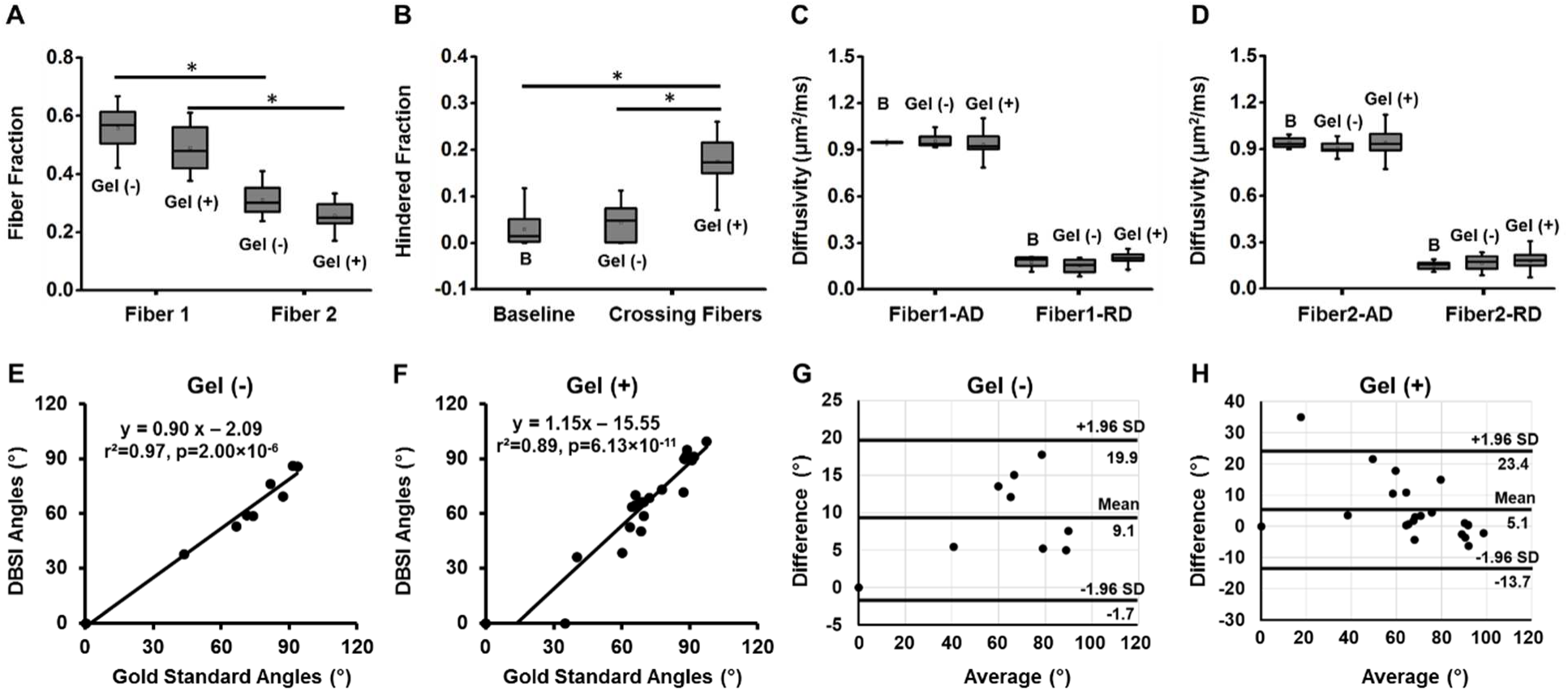
DBSI separated the two crossing fiber complexes and identified individual fiber by their fiber fractions. The DBSI fiber fractions of fiber-1 (juxtaposed fibers) were significantly higher than those of fiber-2 (single fiber) (C, fiber-1 vs. fiber-2, without get: 0.56±0.08 vs. 0.31±0.05; with get: 0.51±0.08 vs. 0.23±0.09). The DBSI hindered fraction of crossing fibers with gel (0.19±0.3) were significantly higher than that of single nerve baseline (0.03±0.04) and crossing fibers without gel (0.04±0.04) (B). DBSI accurately estimated individual diffusivities of both crossing fibers. For fiber 1, the AD and RD of single nerve baseline (AD: 0.95±0.01; RD: 0.17±0.04), crossing fibers without gel (AD: 0.96±0.04; RD: 0.15±0.04) and crossing fibers with gel (AD: 0.94±0.07; RD: 0.20±0.04) were very precise (C). For fiber 2, the AD and RD of single nerve baseline (AD: 0.94±0.03; RD: 0.15±0.02), crossing fibers without gel (AD: 0.91±0.04; RD: 0.16±0.05) and crossing fibers with gel (AD: 0.94±0.08; RD: 0.18±0.05) were also precise (D). DBSI could accurately resolve the fiber crossing angles for conditions without or with gel. The correlation between the DBSI-calculated fiber angles and true crossing angles were significant for those without gel (E, r^2^=0.97, p=2.00×10^−6^) and with gel (F, r^2^=0.89, p=6.13×10^−11^). Bland-Altman analyses indicated the similarity between DBSI estimated fiber angles and true angels for both fibers without gel (G) and with gel (H). Fibers with different amount of gel were summed for analytical analysis. Diffusivity: µm^2^/ms. B = Baseline. P values were adjusted suing Bonferroni correction. *\*P*<0.05.

**Figure 5.**
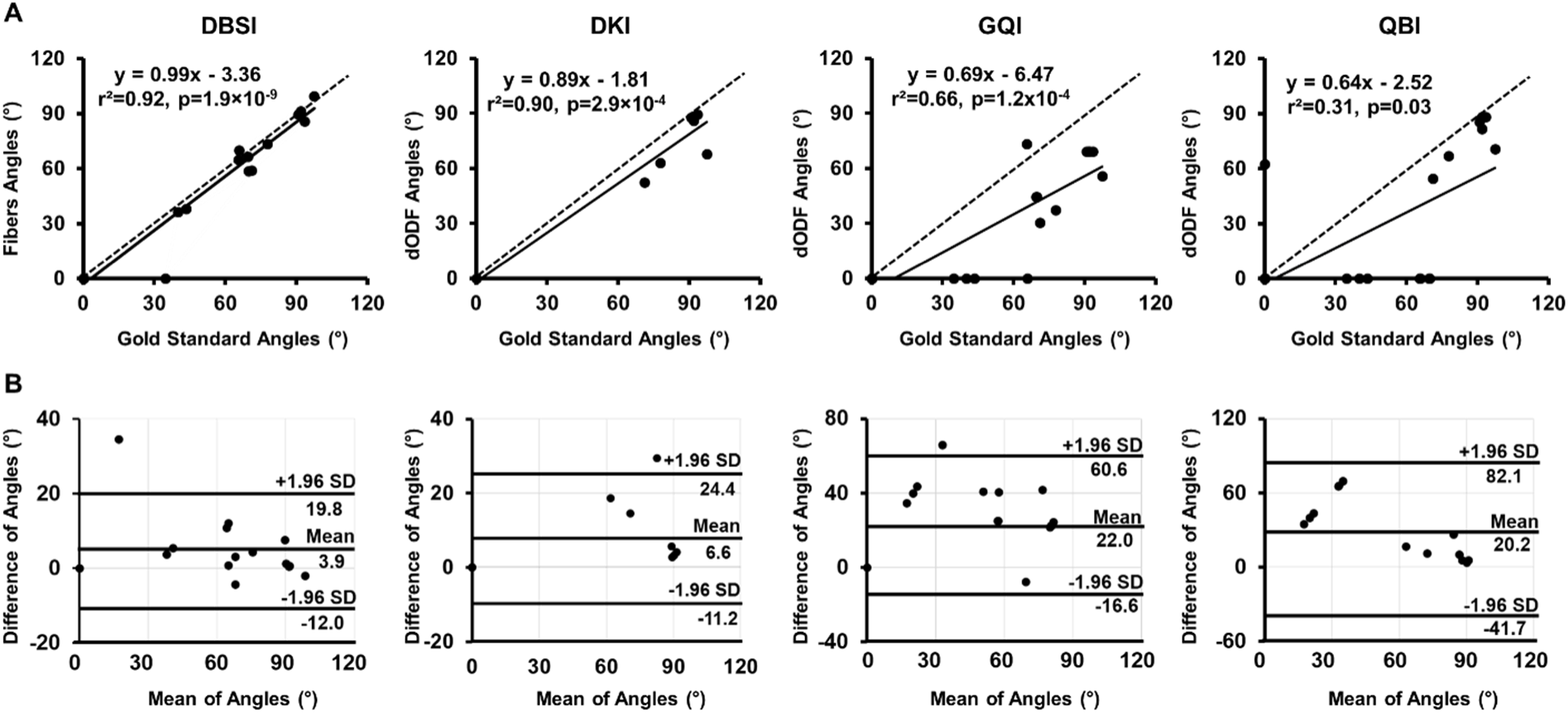
Fiber crossing angles were calculated from DBSI, DKI, QBI, and GQI at various orientations and simulated edematous environments (See Table 1). Pearson correlations (A) and Bland-Altman plots (B) were performed to show the correlations and agreements between dMRI-estimated crossing angles and gold standard angles. Linear lines (dash lines) represent the line of identity for each MRI model (A). Both DBSI- and DKI-estimated crossing angles showed strong correlations with gold standard angles (DBSI: r^2^=0.92, p=1.9×10^−9^; DKI: r^2^=0.90, p=2.9×10^−4^) and small estimated bias (DBSI: bias=3.9°, DKI: bias=6.6°). GQI and QBI showed moderate correlations (GQI: r^2^=0.66, p=1.2×10^−4^; QBI: r^2^=0.31, p=0.33) and larger estimated bias (GQI: bias=22°, QBI: bias=20.2°). Note that DKI detected several crossing fibers with dODF fanning but did not resolve the angles. These datapoints were not included for the above analysis.

**Table 1.** Crossing fiber angles calculated at various orientations and simulated edematous environments from different dMRI methods.

| Phantom Conditions | Gold Standard<br>Angles (°) | dMRI Estimated Angles (°) |  |  |  |
| --- | --- | --- | --- | --- | --- |
|  |  | DBSI | DKI | QBI | GQI |
| fiber a | 0 | 0 | 0 | 0 | 0 |
| fiber b | 0 | 0 | 0 | 0 | 0 |
| fiber c | 0 | 0 | 0 | 0 | 0 |
| fiber d | 0 | 0 | 0 | 0 | 0 |
| fiber e | 0 | 0 | 0 | 0 | 0 |
| fiber f | 0 | 0 | 0 | 62.7 | 0 |
| 40° (f, e) | 43.5 | 38.0 | * | 0 | 0 |
| 40°+coating (d, f+e) | 40.0 | 36.3 | 0 | 0 | 0 |
| 40°+coating (d, f+e) | 34.6 | 0 | 0 | 0 | 0 |
| 60° (d, f+e) | 71.2 | 59.1 | 52.4 | 54.6 | 30.5 |
| 60°+coating (a, b+c) | 65.5 | 64.7 | * | 0 | 73.4 |
| 60°+coating (d, f+e) | 65.8 | 70.1 | * | 0 | 0 |
| 60°+coating+1×gel (a, b+c) | 77.7 | 73.3 | * | 66.8 | 37.4 |
| 60°+coating+1×gel (a, b+c) | 69.7 | 58.8 | 63.1 | 0 | 44.7 |
| 60°+coating+2×gel (a, b+c) | 69.6 | 66.5 | * | 0 | 44.7 |
| 90° (d, f+e) | 93.5 | 85.9 | 89.3 | 88.2 | 69.2 |
| 90°+coating (a, b+c) | 97.4 | 99.5 | 67.8 | 70.9 | 55.8 |
| 90°+coating (d, f+e) | 91.9 | 91.3 | 88.5 | 88.2 | 69.2 |
| 90°+coatin+ 1×gel (a, b+c) | 91.8 | 91.3 | 86.1 | 81.8 | 69.2 |
| 90°+coating+2×gel (a, b+c) | 90.7 | 89.5 | 87.8 | 85.4 | 69.2 |

\* denotes dODF with fanning. dODF with fanning indicates fiber crossing were detected but not resolved.

By accurately detecting and quantifying individual fibers and gel content, DBSI is then able to estimate individual fiber diffusivities with great accuracy (Fig. 4D, E). For both fiber-1 and fiber-2, AD values from crossing fibers without gel (fiber-1: 0.96±0.04 µm^2^/ms; fiber-2: 0.91±0.04) and crossing fibers with gel (fiber-1: 0.94±0.07; fiber-2: 0.94±0.08) were not significantly different (p>0.05) compared with their corresponding baselines (fiber-1: 0.95±0.01; fiber-2: 0.94±0.03). RD values from crossing fibers without gel (fiber-1: 0.15±0.04; fiber-2: 0.16±0.05) and crossing fibers with gel (fiber-1: 0.20±0.04; fiber-2: 0.18±0.05) were also not significantly different (p>0.05) compared to baseline (fiber-1: 0.17±0.04; fiber-2: 0.15±0.02). From the above results, DBSI, unlike DTI, successfully calculated AD and RD regardless of crossing angles and extent of edema. DBSI also accurately resolved the fiber crossing angles with or without gel. The DBSI-calculated fiber angles correlated with gold standard crossing angles, without gel (Fig. 4E, r^2^=0.97, p=2.00×10^−8^) and with gel (Fig. 4F, r^2^=0.89, p=6.13×10^−11^). Bland-Altman analyses also indicated close agreements between DBSI-estimated fiber angles and gold standard angles with bias (average of difference) of 9.1° (without gel, Fig. 4H) and 5.1° (with gel, Fig. 4I), respectively. This indicated DBSI is able to accurately estimate crossing angles with small variances under various edematous conditions.

It is important to note that DKI, GQI and QBI, as detailed in the introduction section of this paper, are not designed to estimate the diffusivity of individual fibers while simultaneously resolving crossing angles. Thus, none of these methods was performed to estimate the impact of fiber crossing and edema on diffusivity.

### 3.3 Comparing DBSI and dODF (DKI, GQI and QBI) detected and esitmated crossing angles

The dODF is used with various modifications by DKI, GQI, and QBI to resolve fiber crossing. To determine whether these models could accurately resolve crossing angles, we compared crossing angles estimated by DKI, GQI, QBI, and DBSI to gold standard angles at approximately 90°, 60°, and 40° in different gel-mimicked edematous conditions (Table 1)

Overall, DBSI correctly detected more single/crossing fibers than other dMRI methods. DBSI correctly identified 19/20 (19 out of 20) samples, compared with 18/20 samples from DKI, with 16/20 samples and 12/20 samples from QBI (Table 1). Additionally, DBSI exhibited better crossing angle calculations with respect to the gold standard angles (r^2^=0.92, p=1.9×10^−9^, bias=3.9°). DKI, GQI, and QBI all estimated crossing angles but were not as close to the gold standard angles (r^2^=0.90, p=2.9×10^−4^, bias=6.6°; r^2^=0.66, p=1.2×10^−6^, bias=22°; and r^2^=0.31, p=0.03, bias=20.2°, respectively). Note that DKI identified five crossing fibers but did not resolve the angles, thus they were not included in the above analyses.

For 90-degree crossing angle with/without gel, DKI, QBI, and DBSI all performed well (Table 1). GQI resolved all the 90° crossing angles but with deviations of ~20°. In nerve assembly of 60° crossing, DKI, QBI and GQI partially resolved the crossing angles but with less accuracy than DBSI. DBSI resolved all 60° angles, and the estimates did not deviate from the gold standard angle as more gel was added. Importantly, DKI detected all six crossing fibers but failed to resolve the angles in four instances, all of which were under gel conditions. DKI reconstructed the dODF with fanning that successfully identified the fiber dispersion but was unable to estimate the crossing angles (Fig. S1). Compared to DKI, QBI failed to detect four out of six crossing. GQI detected and calculated five of the six crossing angles but with substantial underestimates in comparison to the gold standard. At 40° crossing, DBSI was able to calculate two crossing fibers with close estimates of angles but failed in one attempt. None of the other methods resolved these small angles. In particular, DKI showed dODF with fanning in one case but failed in two other instances. Interestingly, one single fiber was mischaracterized by QBI as having a crossing angle of 62.7°. In sum, small crossing angles and gel-mimicked edema prevented dODF from detecting and/or estimating crossing angles. In contrast, DBSI by adopting multi-tensor stragedy, successfully detected and estimated various crossing angles with different get consitions.

Although NODDI was not designed to calculate fiber crossing angles, it calculated orientation dispersion index to quantify the bending and fanning of axons. Here we also compared NODDI derived ODI with gold standard angles to see whether ODI could reflect different degrees of neurite dispersion. In general, ODI indicated strong positive correlation with gold standard crossing angles for dataset with single fibers (Fig. S2A, r^2^=0.60, p=0.0007) and dataset without single fibers (Fig. S2B, r^2^=0.75, p=5.8×10^−5^). However, single nerves indicated close ODI values to 40° crossing fibers (Table 1, 0.83±0.02 vs. 0.82±0.01), which was unexpected (Table S1).

### 3.4 DBSI accurately characterizes brain tumor and surrounding white matter tissue via white matter tractography

MR images and whole brain white matter tractography maps from a patient with metastatic lung carcinoma were collected (Fig. 6). Gd-enhanced T1W images showed heterogeneous enhancements in the right frontal lobe (Fig. 6A, yellow arrowhead), indicating presence of solid tumor. FLAIR-T2W image showed peritumoral edema (hyper-intense signals) around Gd-enhanced region (Fig. 6B, green arrowhead). From direction-encoded DBSI tractography map, we detected healthy white matter tract distribution within the peritumoral region with vasogenic edema (Fig. 6C green arrow, 6E), which was represented as vasogenic edema with no heathy white matter tracts by DTI-derived tractography map (Fig. 6D green arrow, 6F). DBSI identified white matter tracts amidst the surrounding edema (Fig. 6E) while DTI failed to detect these tracts (Fig. 6F). Isotropic hindered diffusion fraction was encoded onto the DBSI tractography tracts to highlight the extent of vasogenic edema (Fig. 6E, square). DTI-derived ADC values were also encoded onto DTI tractography but did not reveal presence of healthy underlying white matter tracts (Fig. 6F).

**Figure 6.**
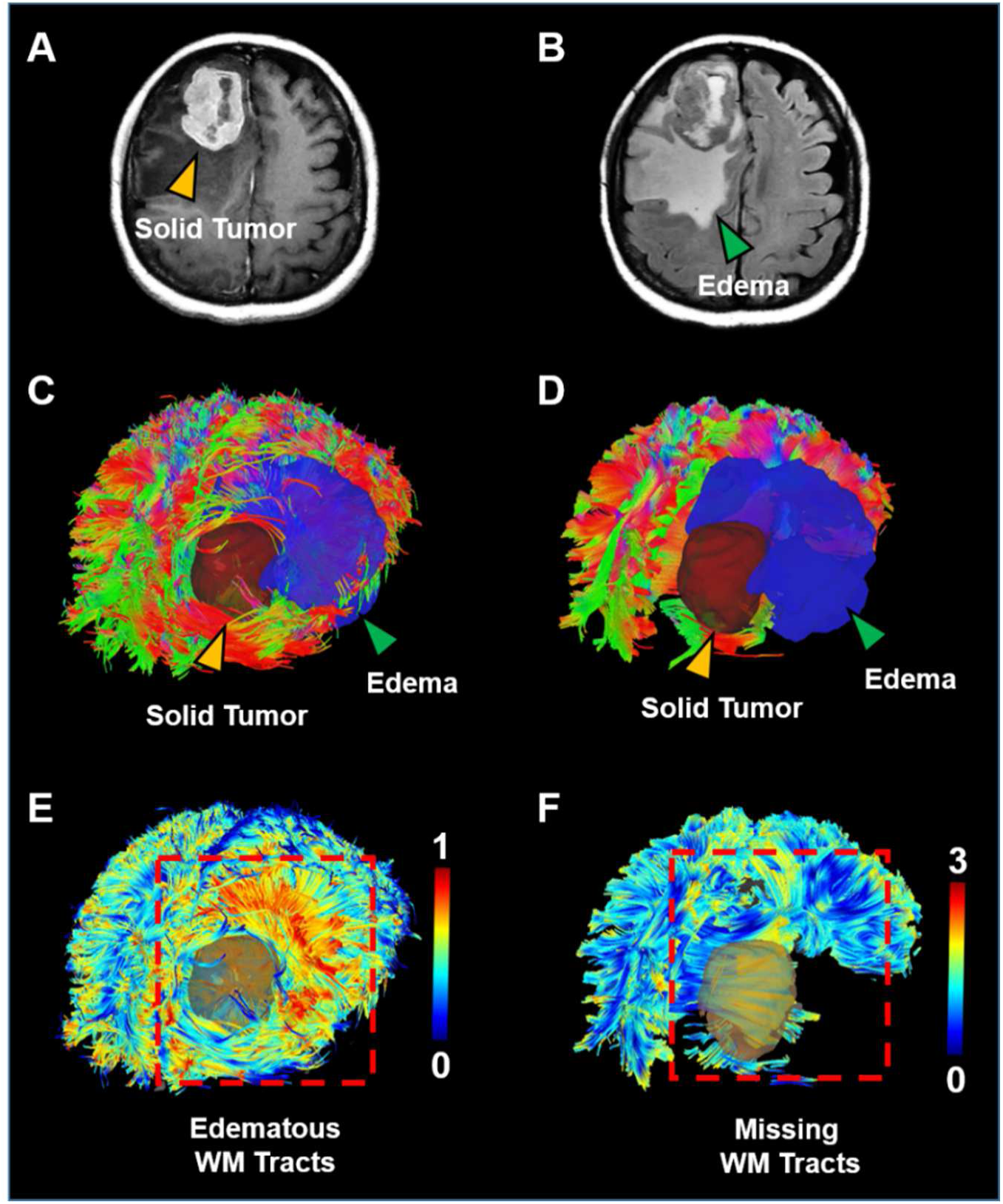
MRI images and whole brain white matter tractography maps from a metastatic lung carcinoma patient. Gd-enhanced T1W images showed heterogeneous enhancements in the right frontal lobe region (A, yellow arrowhead), which indicates solid tumor area. FLAIR-T2W image showed hyper intense signals around Gd-enhanced region (B, green arrowhead), indicating peritumoral edema. DBSI successfully recovered white matter tracts within the surrounding edematous area (C, green arrow), while DTI failed to detect white matter tracts in the peritumoral edematous area on direction-encoded map (D). DBSI-hindered-fraction-encoded tractography map showed higher hindered fraction for edematous tracts (E), which revealed the peritumoral edema grew and spread along the white matter tracts (E). DTI-ADC-encoded map identified the tumor and surrounding edematous area (D) but failed to identify underlying healthy white matter tracts (F). Direction-encoded white matter map is oriented as follows: red = left to right; green = anterior to posterior; blue = superior to inferior.

## 4 DISCUSSION

We investigated DBSI’s efficacy for simultaneously estimating individual fiber diffusivity and crossing fiber angles under conditions of gel-mimicked edema. We compared DBSI’s results with those of DTI and other advanced diffusion MRI techniques. Mouse trigeminal nerves were aligned in a 1:2 ratio at three different crossing fiber angles and three different simulated environments of edema. Nerve diffusivity was estimated by both DTI and DBSI and compared to baseline diffusivity values for each nerve. DBSI more accurately estimated AD and RD with less variation than DTI estimations. Compared with single nerve control, mean DTI AD decreased by 13%, 16%, and 22% for 40, 60, and 90 degree angles, respectively; DTI RD increased by 80% and 120% for 60 degree and 90 degree angles, respectively; DTI FA decreased by 3%, 38% and 53% for 40, 60, and 90 degree angles, respectively (Fig. 3). DTI, therefore, is not able to accurately recover diffusivity and FA from individual crossing-fiber.

DBSI distinguished the individual fibers from the two-fiber complex and accurately estimated the respective AD and RD for both the single nerve and two-nerve complex (Fig. 4). There were no significant differences of AD and RD between the gel-coated nerves with respect to single nerve baseline. These results suggest that DBSI-derived AD and RD are representative assessments of axon injury and demyelination in the presence of coexisting pathologies such as vasogenic edema and inflammatory infiltration. DBSI separates anisotropic diffusion components from the isotropic diffusion, enabling DBSI to correctly distinguish individual nerves and accurately quantify their diffusivities. The fiber fraction of each nerve is calculated from the signal fraction of each anisotropic diffusion component.

Fiber crossing is ubiquitous within the human brain; resolving crossing fibers is needed not only to obtain accurate axial and radial diffusivity measures, but to perform anatomical connectivity studies with precision.^32^ Although dODF has been widely adopted by many diffusion MRI methods to determine fiber orientations and resolve crossing fibers, their accuracy could be limited by the blurred contour of dODF.^33^ As demonstrated in Results section above, shortcomings of dODF may result in failure to resolve crossing angles, especially acute angles, in edematous environments. To overcome the limitations of dODF, fiber ODF (fODF) has been proposed and calculated using spherical deconvolution by several dMRI methods, such as constrained spherical deconvolution (CSD)^34^ and ball-and-sticks model,^35^ to measure the orientation distribution of fiber volume fractions.^36,37^ In general, fODF could provide a sharper contour and achieve better angular resolution than dODF. Despite fODF has been demonstrated to be sensitive to detect crossing fibers, studies have cast doubts on its specificity.^38,39^ Especially, fODF regularly possesses baseline fluctuations that could result in false peaks and increase false detections.^33^ In addition to the baseline fluctuation, the employmemt of L2 regularization also induces blurring to fODF, which could undermine the accurate estimation of crossing fibres.^33^

In contrast to dODF and fODF, DBSI adopts a strategy of modeling DWI data as a linear combination of multiple discrete anisotropic tensors and a spectrum of isotropic diffusion tensors.^17^ Fiber orientations were determined from discrete diffusion basis sets of variable diffusivities while coexisting cells, CSF, and edema was modeled by a spectrum of isotropic diffusion tensors, enabling a non-confounded analysis of the crossing angle.^17^ Our study revealed that while DBSI, QBI, GQI, and DKI could resolve crossing angles of ~90° in the four different simulated pathologies, DBSI did so with better accuracy and precision. Furthermore, DBSI accurately and consistently estimated fiber crossings at smaller angles of ~60° and ~40°, which outferformed other dMRI methods.

White matter tractography has been widely utilized in invasive surgical procedures to minimize healthy tissue resection and maximize diseased tissue resection.^40,41^ DTI-based tractography, however, has proven inadequate in edematous areas due to the reduction of FA by partial volume effects.^42–44^ In particular, decreases in FA can lead to prematurely terminated DTI tractography representations of perilesional nerve tracts.^45^ Our results are consistent with literature reports that DTI tractography failed to detect white matter tracts in the peritumoral regions of edema. However, our results demonstrate that DBSI simultaneously avoids structural confounds such as edema/CSF contamination, estimates accurate crossing fiber angles, and accurately derives AD and RD for resolved fibers. Thus, DBSI is a more accurate and robust diffusion MRI model for evaluating nerve crossing fibers in edematous environments. Further, we showed that DBSI accurately detects white matter tracts in peritumoral edematous areas in a patient with brain tumor (Fig. 6). The high hindered fractions of the edematous white matter tracts demonstrate that DBSI is able to detect, distinguish, and quantify fibers within the edematous peritumoral region in a patient with brain tumor. Meanwhile, DTI is unable to identify healthy white matter tracts in the peritumoral region, suggesting that DBSI’s ability to resolve crossing fibers at sub-voxel levels provides a more robust model for deterministic tractography model than DTI.

There are several limitations of this study. Firstly, the number of trigeminal nerves used in the experiments was limited. The mouse trigeminal nerves are very delicate. Some nerves were irreversibly damaged during the extraction and slide preparation process or from the gelling process. The experimental design requires repeated usage of nerves, which required discarding of even slightly damaged nerves, lowering the sample size of the study. The low sample size prohibited comparison of nerve diffusivities at all three gel environments. Rather, they were averaged into a “gel+” environment for DBSI and DTI comparisons. This prohibited diffusivity and crossing angle analyses at increasing levels of edema, which we intended to serve as a proxy of increasing severity of disease. Although combining the various levels of added gel into one group limited the depth of analyses that we could perform, we were able to demonstrate marked differences in diffusivity estimations, and, for the first time, differences in various diffusion MRI models’ abilities to resolve crossing fiber angles. Secondly, the performance of DKI dODF in resolving crossing angles could be compromised as DKI analyses were performed based on the partial dataset from LEMONADE. In future studies, we will address this by adopting the recommended diffusion scheme for each diffusion MRI model. Thirdly, the experimental design of this phantom study that used fixed mouse trigeminal nerves and agarose gel could possibly lead to the outperformances of DBSI over dODFs by DKI, QBI and GQI, as the underlying assumptions may be better justified for DBSI in these phantoms. The potential advantage of dODFs is that they are less dependent on sample-specific assumptions. In future study, it would be crucial to compare DBSI, fODF and dODF in resoling crossing fibers under various conditions and samples, e.g., human brain study.

## 5 CONCLUSIONS

Our results demonstrate that DBSI distinguishes nerve bundles and calculates *ex vivo* axonal diffusivities and crossing fiber angles precisely and accurately. Preliminary investigation into DBSI’s clinical efficacy for differentiating fibers from surrounding edematous conditions demonstrates that DBSI differentiates brain tumor cells from surrounding edematous tissues *in vivo*. Our small sample size, however, necessitates further study into this comparison to make a generalizable determination. In summary, DBSI’s ability to quantify multiple sub-voxel diffusion components confers an unprecedented ability for simultaneously providing multiple specific pathological biomarkers in CNS diseases and brain tumor. Its longitudinal use in patients with CNS diseases like multiple sclerosis and brain tumor could potentially improve the understanding and treatment stratification of the disease.

## Author Contributions

S.K.S., Z.Y., and S.G. conceived and designed the study. Z.Y. and S.G. performed the experiments. Z.Y., S.G., P.S., G.R.G., S.M.M., F.C.Y. and H.M analyzed the experimental data. Z.Y., S.G. and S.K.S. wrote the manuscript, and was assisted by J.J., and W.Y.C.. All authors have reviewed and approved the final version of the manuscript.

