## Supplementary Figure for "The Impact of Edema and Fiber Crossing on Diffusion MRI Metrics: DBSI vs. Diffusion ODF"

### Supplements

### Supplementary Figures

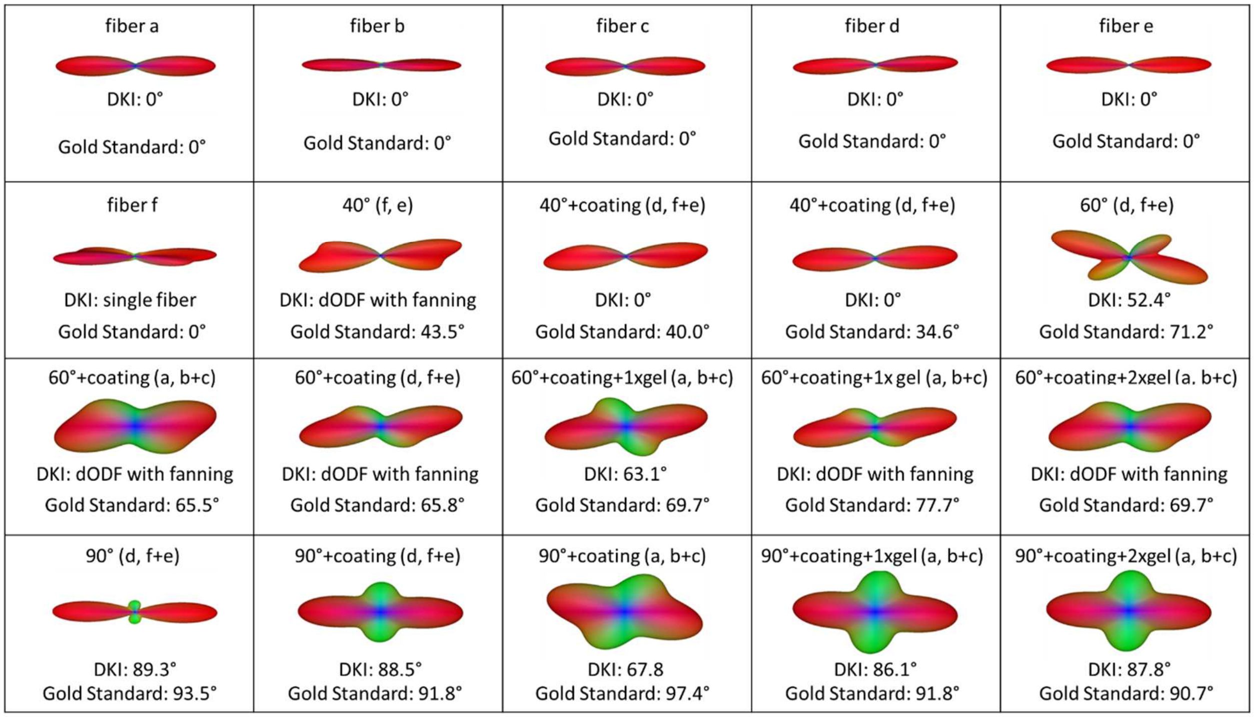


**Figure S1.** DKI dODF reconstruction and estimated fiber crossing angles compared with gold standard angles under various edematous environments and crossing angles.

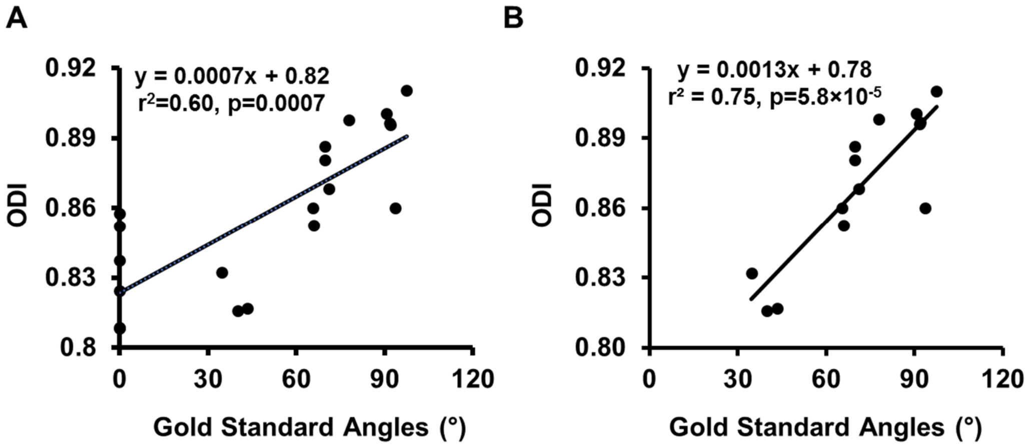


**Figure S2.** Correlaitons between NODDI-derived ODI and gold standard angles for dataset with single fiber (A) and without single fiber (B).

**Table S1.** NODDI derived ODI calculated at various fiber crossing angles and simulated edematous environments .

| Phantom Conditions | Gold Standard Angle (°) | NODDI ODI |
|---|---|---|
| fiber a | 0 | 0.857 |
| fiber b | 0 | 0.824 |
| fiber c | 0 | 0.852 |
| fiber d | 0 | 0.808 |
| fiber e | 0 | 0.809 |
| fiber f | 0 | 0.837 |
| 40° (f, e) | 43.5 | 0.817 |
| 40°+coating (d, f+e) | 40.0 | 0.832 |
| 40°+coating (d, f+e) | 34.6 | 0.816 |
| 60° (d, f+e) | 71.2 | 0.868 |
| 60°+coating (a, b+c) | 65.5 | 0.860 |
| 60°+coating (d, f+e) | 65.8 | 0.852 |
| 60°+coating+1×gel (a, b+c) | 77.7 | 0.898 |
| 60°+coating+1×gel (a, b+c) | 69.7 | 0.881 |
| 60°+coating+2×gel (a, b+c) | 69.6 | 0.886 |
| 90° (d, f+e) | 93.5 | 0.860 |
| 90°+coating (a, b+c) | 97.4 | 0.910 |
| 90°+coating (d, f+e) | 91.9 | 0.897 |
| 90°+coatin+ 1×gel (a, b+c) | 91.8 | 0.896 |
| 90°+coating+2×gel (a, b+c) | 90.7 | 0.900 |
